# Evolution and Genetic Diversity of SARSCoV-2 in Africa Using Whole Genome Sequences

**DOI:** 10.1101/2020.07.27.222901

**Authors:** Babatunde Olarenwaju Motayo, Olukunle Oluwapamilerin Oluwasemowo, Paul Akiniyi Akinduti, Babatunde Adebiyi Olusola, Olumide T Aerege, Adedayo Omotayo Faneye

## Abstract

The ongoing SARSCoV-2 pandemic was introduced into Africa on 14^th^ February 2020 and has rapidly spread across the continent causing severe public health crisis and mortality. We investigated the genetic diversity and evolution of this virus during the early outbreak months using whole genome sequences. We performed; recombination analysis against closely related CoV, Bayesian time scaled phylogeny and investigated spike protein amino acid mutations. Results from our analysis showed recombination signals between the AfrSARSCoV-2 sequences and reference sequences within the N and S genes. The evolutionary rate of the AfrSARSCoV-2 was 4.133 × 10^−4^ high posterior density HPD (4.132 × 10^−4^ to 4.134 × 10^−4^) substitutions/site/year. The time to most recent common ancestor TMRCA of the African strains was December 7^th^ 2019. The AfrSARCoV-2 sequences diversified into two lineages A and B with B being more diverse with multiple sub-lineages confirmed by both maximum clade credibility MCC tree and PANGOLIN software. There was a high prevalence of the D614-G spike protein amino acid mutation (82.61%) among the African strains. Our study has revealed a rapidly diversifying viral population with the G614 spike protein variant dominating, we advocate for up scaling NGS sequencing platforms across Africa to enhance surveillance and aid control effort of SARSCoV-2 in Africa.

## INTRODUCTION

Towards the end of December 2018, Chinese authorities through the World Health Organization office in China made known of a new pathogen responsible for a series of pneumonia associated infections in Wuhan, Hubei province (WHO 2020). The pathogen was later identified to be a novel coronavirus closely related to the severe acute respiratory syndrome virus (SARS), with a possible bat origin (Zhou et al, 2020). The world health organization named the disease COVID-19 (Chan et al, 2020), and later declared it a pandemic on 11^th^ March 2020 prompting concerted efforts towards prevention and control worldwide (WHO 2020). On Febuary 11^th^ 2020 the international committee on the taxonomy of viruses (ICTV) adopted the name SARS-CoV-2 following the report of their coronavirus working group (CSG, 2020). The virus has been placed in the subgenera *sarbecovirus*, genus *betacoronavirus*, subfamily coronavirinea, family Coronaviridea (de Groot et al, 2013; Gorbalenya et al, 2020).

Coronaviruses are enveloped viruses containing a single-stranded positive sense RNA genome with a size of between 26kb to 32kb (Masters and Pearlman 2013). They are responsible for a host of human and animal infections. The *betacoronaviruses* host the most medically important species contains several human coronaviruses such as HuCoVOC43, HuCoVHKu13. The severe acute respiratory syndrome coronavirus SARSCoV and the Middle East respiratory syndrome coronavirus MERS are also members of this group, both have been shown to be pathogens of high consequence that caused large scale epidemics and have been shown to be of zoonotic origin (Lau et al, 2005; Zaki et al, 2012). Genomic and structural analyses have revealed that SARSCoV-2 contains four structural proteins and several non structural proteins (Chen et al, 2020; Lu et al, 2020). The spike protein is the major antigenic protein responsible for initiating infection, via attachment of its receptor binding domain (RBD) to the SARSCoV/SARSCoV-2 receptor angiotensin converting enzyme 2 ACE 2 (Donelli et al, 2004; Monteil et al, 2020).

Globally, there have been 5,226,268 confirmed SARSCoV-2 cases globally, with 335,218 deaths as at 21^st^ of May 2020 (ECDC 2020). The coronavirus pandemic began in Egypt Africa on the 14^th^ Febuary 2020 with an Italian who returned into the country (WHO 2020b). As at 21^st^ of May there have been 95,332 cases in Africa, with 2,995 deaths and 35,519 recoveries covering fifty four countries in Africa with South Africa having the highest number of cases of 18,003 (WHO 2020c). Several reports have traced the evolutionary origins of SARSCoV-2 to SARSrCoV from bats (Zhou et al, 2020) and Pangolins (Lam et al 2020).

Phlogenetic analysis has shown that the virus has diversified through the duration of the pandemic into three major lineages A and B with several sub-lineage diversifications (Rambault et al 2020). Majority of the reports were generated using genome sequences of SARSCoV-2 from America, Europe and Asia (Rambault et al, 2020). There has been paucity of data on the genetic evolution of SARSCoV-2 sequences from Africa, despite the increasing number of genome sequence submissions into the GISAID database from Africa; there were 97 whole genome sequences available in the GISAID database as at 24^th^ April 2020. This gap in knowledge prompted the conceptualization of this study. This study was designed to determine to the genetic diversity and evolutionary history of genome sequences of SARSCoV-2 isolated in Africa.

## MATERIALS AND METHODS

### DATA CURATION

Full genome sequences with high coverage were downloaded from the Global initiative for sharing of Avian Influenza data GISAID database. As at 24^th^ April there were 97 full genome sequences from Africa available in the GISAID database we downloaded all of them excluding genomes with low coverage. A total of 69 high coverage genomes were eventually selected from the African sequences, along with these 151 high quality full genome sequences were also downloaded from three continents America (USA), Asia (China and South Korea) and Europe (England, Italy and Germany). Three different datasets were then generated from these sequences, the first dataset consisted of high coverage full genome sequences from Africa, along with the SARSCoV2 reference genome sequence from Wuhan, China, Bat and Pangolin SARS related reference sequences and SARSCoV reference sequence. The second dataset consisted of complete genome sequences from Africa, America, Asia and Europe, while the third dataset consisted of complete spike protein (S) gene sequences from Africa, Bat and Pangolin SARS related reference S gene sequences.

### PHYLOGENETIC ANALYSIS

Whole genome sequences downloaded from the GISAID database were aligned using MAFFTv7.222 (FF-NS-2 algorithm) following default settings (Katoh et al, 2019). Maximum likelihood phylogenetic analysis was performed using the general time reversible nucleotide substitution model with gamma distributed rate variation GTR-Γ(Yang et al, 1994) with 1000 bootstrap replicates using IQ-TREE software (Nguyen et al 2015). Lineage assignments for the SARSCoV-2 sequences were conducted using the Phylogenetic Assignment of Named Global Outbreak LINeages tool (PANGOLIN), available at http://github.com/hCoV-2019/pangolin (O’Toole and McCrone 2020).

### RECOMBINATION ANALYSIS

We analyzed potential recombination events using the recombination detection program RPD software (Martin et al, 2015). The analysis was conducted on whole genome sequences of identified lineages among the African isolates, using RDP, bootscan analysis, GENECOV, Chimera, SISCAN, 3SEQ, and maximum chisquare methods. A putative recombination event was passed only if three of the above mentioned methods gave a positive recombination signal (Liu et al, 2010).

### EVOLUTIONARY AND TIME SCALED PHYLOGENETIC ANALYSIS

Temporal clock signal was analyzed among the aligned sequences using TempEst version 1.5 (Rambault et al, 2016). The root-to-tip divergence and sampling dates supported the use of molecular clock analysis in this study. Phylogenetic trees were generated by Bayesian inference through Markov chain Monte Carlo (MCMC), implemented in BEAST version 1.10.4 (Suchard et al, 2016). We partitioned the coding genes into first+second and third codon positions and applied a separate Hasegawa-Kishino-Yano (HKY+G) substitution model with gamma-distributed rate heterogeneity among sites to each partition (Hasaegawa et al, 1985). The relaxed clock with Gausian Markov Random Field Skyride plot (GMRF) coalescent prior was selected for the final analysis, after running different models and comparing them using Bayes factor with marginal likelihood estimated using the path sampling and stepping stone methods implemented in BEAST version 1.10.4 (Suchard et al, 2016). One hundred million MCMC chains were run with10% burn in. Results were then visualized with Tracer version 1.8. (http://tree.bio.ed.ac.uk/software/tracer/), all effective sampling size ESS values were >200 indicating sufficient sampling. Bayesian skyride analysis was carried out to visualize the epidemic evolutionary history using Tracer v 1.8.

### ANALYSIS OF SARSCoV-2 SPIKE PROTEIN

Complete S protein gene sequence of AfrSARSCoV-2 was aligned along with RaTG13 BtCoV and Pangolin SARSrCoV sequences using MAFFT (Katoh et al, 2015). The alignment was then edited and visualized using BioEdit software.

## RESULTS AND DISCUSSION

The current global SARSCoV-2 pandemic, otherwise known as COVID-19 began on the African continent from a European returnee in Egypt on February 17^th^ 2020 (WHO 2020). It has since spread to virtually all the countries within the African region. This study was based on sequences generated during the early phase of the pandemic in Africa precisely between, February 2020 and April 2020. Sixty nine high coverage full genome sequences from six African countries, namely Algeria (3), Senegal (20), Democratic republic of Congo DRC (35), Nigeria (1), Ghana (6) and South Africa (4) were analyzed. Phylogenetic analysis of the African sequences showed clustering within the *sarbecovirus* sub-genus forming a sub-cluster with SARSr CoV and PCoV (Figure 1) as previously reported by several workers(Zhao et al, 2020; Lam et al, 2020; Zhang et al, 2020). The root to tip regression analysis showed a not so strong signal with a correlation of coefficient of 0.995 and R^2^ = 0.991 (Supplementary Figure 1).

**FIGURE 1.**
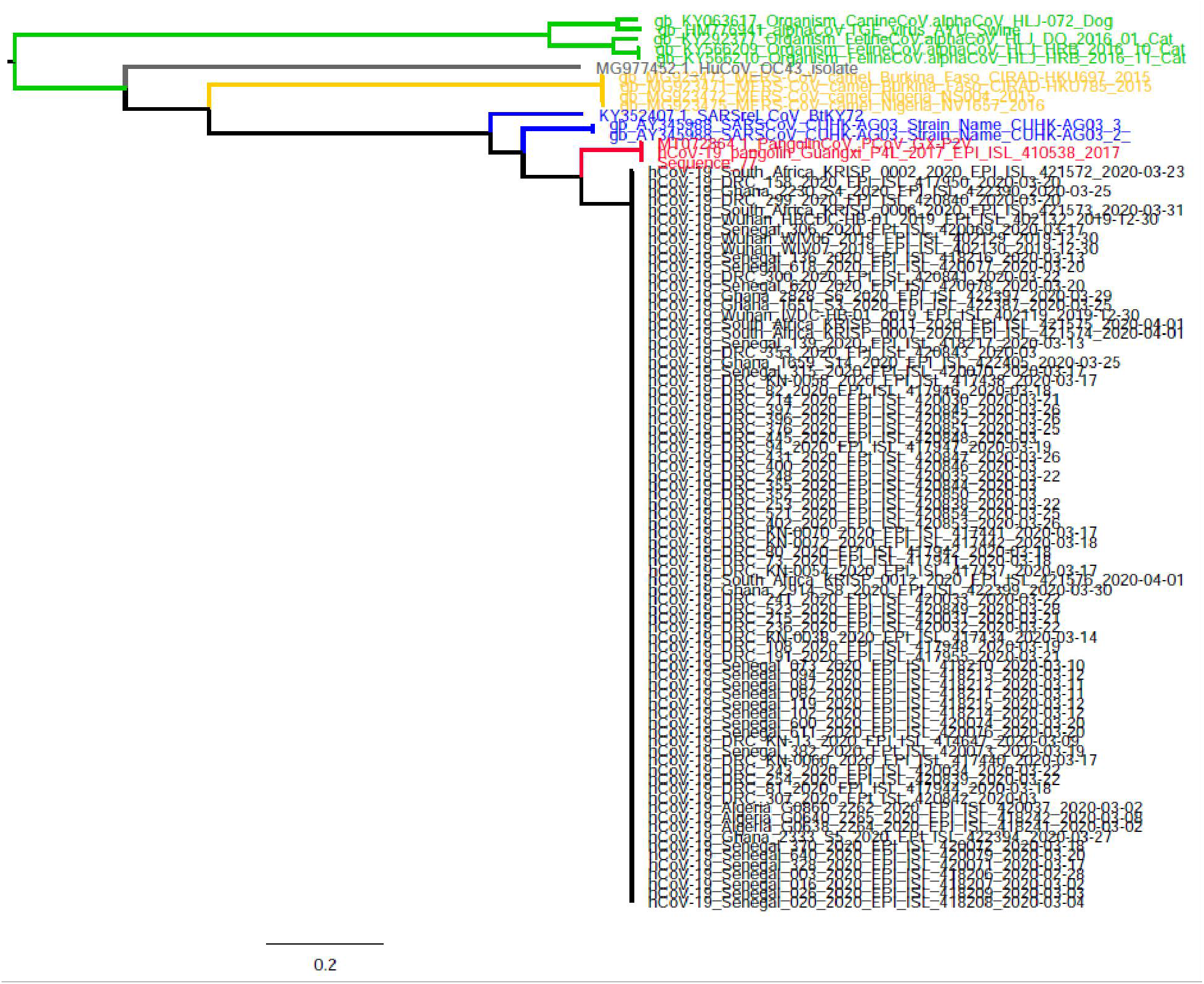
Maximum Likelihood phylogenetic tree of AfrSARSCoV2 full genome sequences along with Pangolin SARS-like, Bat SARS-like Cov, SARSCoV, MERSCoV and alpha CoV full genome sequences.

Results of recombination analysis of the African SARSCoV-2 (AfrSARSCoV-2) sequences against references whole genome sequences of SARS, Recombination signals were observed between the African SARSCoV-2 sequences and reference sequence (Major recombinant hCoV-19 Pangolin/Guangu P4L/2017; Minor parent hCoV-19 B batYunan/RaTG13) between the RdRP and S gene regions (Figure 2). This result is consistent with a previous report from Saudi Arabia which investigated the recombination between SARSCoV-2 and closely related viruses such as SARSCoV and MERS (Nour et al, 2020). Evolutionary rate for the AfrSARSCoV-2 isolates during the period under study was 4.133 × 10^−4^ substitutions/site/year, high posterior density interval HPD (4.132 × 10-4 to 4.134 × 10-4). This is slightly higher than that of an earlier report from early outbreak strains from China with a rate of 3.345 × 10-4 (Li et al, 2020), it is however lower than the calculated global SARSCoV-2 evolutionary rate estimated to be 8.0 × 10-4 reported by Nexstrain (www.nextstrain.org/ncov/global). The MCC tree of the African SARSCoV-2 sequences shows that they have evolved into two major lineages A and B with lineage B being more diverse. Majority of the African SARSCoV sequences clustered within lineage B, while three Ghanaian, three Congolese, and four Senegalese strains clustered along with the reference Chinese and South Korean strains within lineage A (Figure 3). The MCC tree for the dataset containing global reference sequences also showed a similar topology with that of the African tree, the tree was distributed into two major lineages A and B, with lineage B further diversifying into about four sub-lineages, while lineage A seemed to evolve into only two sub-lineages (Figure 4). The AfrSARSCoV-2 strains were intermixed with the global sequences within both lineages, lineage B consisted mainly of strains from Germany, England, Italy and USA, intermixed with African strains; while lineage A consisted mainly of strains from South Korea and China with a few African strains from Senegal, Ghana and DRC. The result of the genotype analysis using the genotyping tool PANGOLIN was largely in conformity with observed phylogenetic analysis. Figure 5 shows a summary of the lineage distribution of the isolates by country of origin using the PANGOLIN genotyping tool. The complete distribution of the strains according to lineage and country is shown in supplementary Table 2. From the analysis with PANGOLIN, lineage B.1 was the most commonly encountered and the most widely distributed, consisting of 93 sequences from Seven countries, followed by lineage B.2 and genotype B. Lineage A had 15 positive sequences from six countries. Majority of the sequences recorded high bootstrap values with over 70% of the sequences recording a bootstrap value of above 80%. This shows that the PANGOLIN is a reliable tool with a broad scope of functions including a user friendly and interactive representation of phylogenetic clustering of the identified sub-lineages and lineages by means of graphical images of the trees generated using virtually all available SARCoV-2 sequences available on GISAID platform as reference. The genotyping tool was recently introduced several reports have utilized it in predicting lineage assignments accurately (Xaiveir et al, 2020). The time to most recent common ancestor TMRCA of the African SARSCoV-2 strains was December 7^th^ 2019 (November 12^th^ 2019-December 29^th^ 2019), while the TMRCA of all the sequences under analysis was 14^th^ October 2019 (July 27^th^ 2019-December 17^th^ 2019). Our TMRCA was lower than a similar study which reported a TMRCA of 14^th^ October 2019 among global isolates including Chinese isolates (Li et al, 2020), but was slightly higher than another recent study investigating the evolutionary dynamics of the ongoing SARSCoV-2 epidemic in Brazil which reported a TMRCA of 10^th^ February 2020 (Xaiveir et al, 2020). The epidemic history of the ongoing outbreak was investigated using the Bayesian Skyline Plot BSP. The BSP showed a steady increase in viral population as the outbreak progressed under the study period (Figure 6). This observation is expected as viral sequence population is supposed to increase as the infection spreads. A major limitation was the rather small number of sequences analyzed and very short study duration; therefore our results may not reflect the exact viral population dynamic of the outbreak in Africa.

**FIGURE 2.**
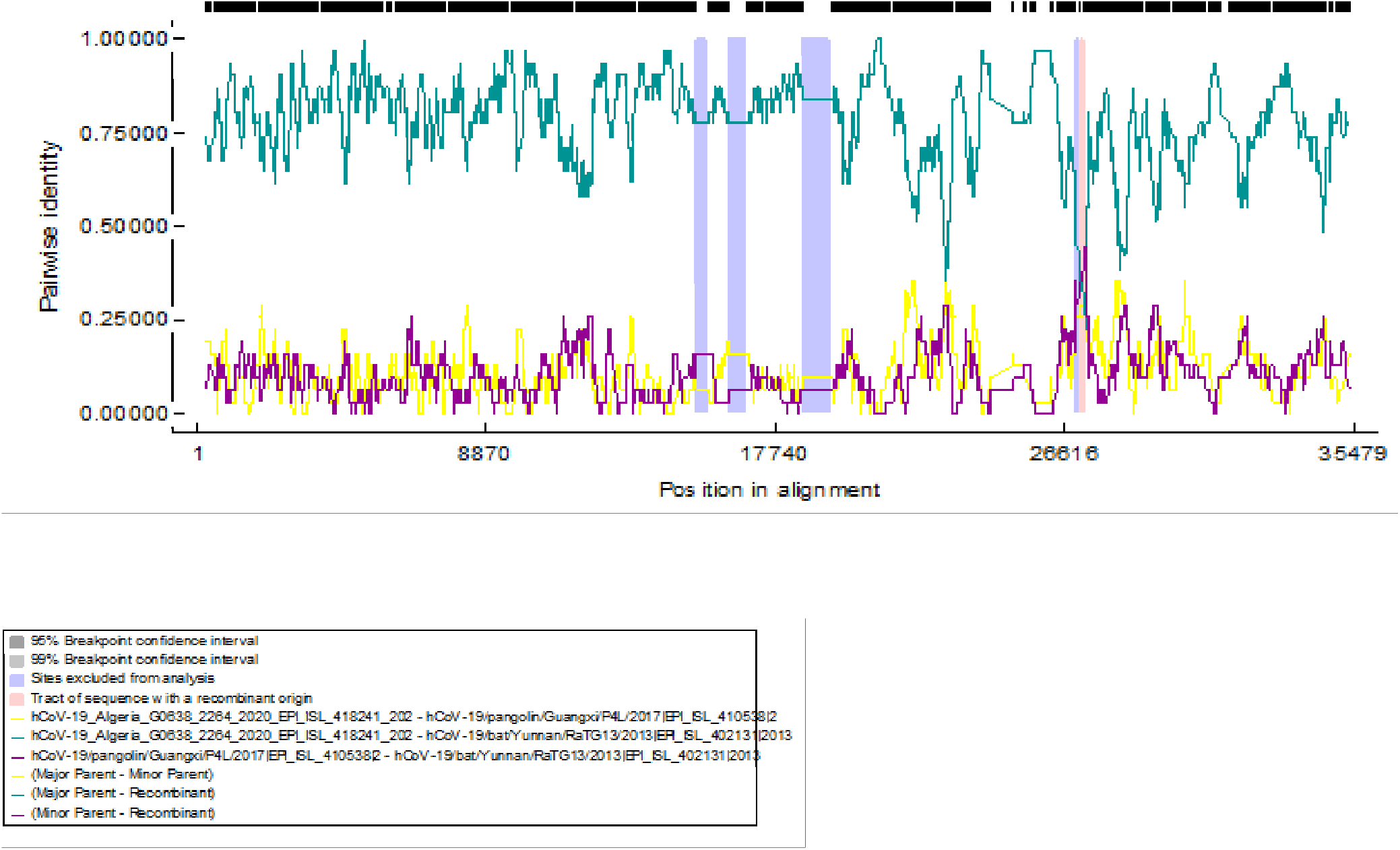
Boot scan plot of complete genome sequences of AfrSARSCoV-2 sequences analysed with the RDP recombination software. The legend shows the identity of the sequences scanned within the plot; the light blue bars indicate the portions of the genome with recombinant signals in reference to the Major and minor recombinant parent sequences.

**FIGURE 3.**
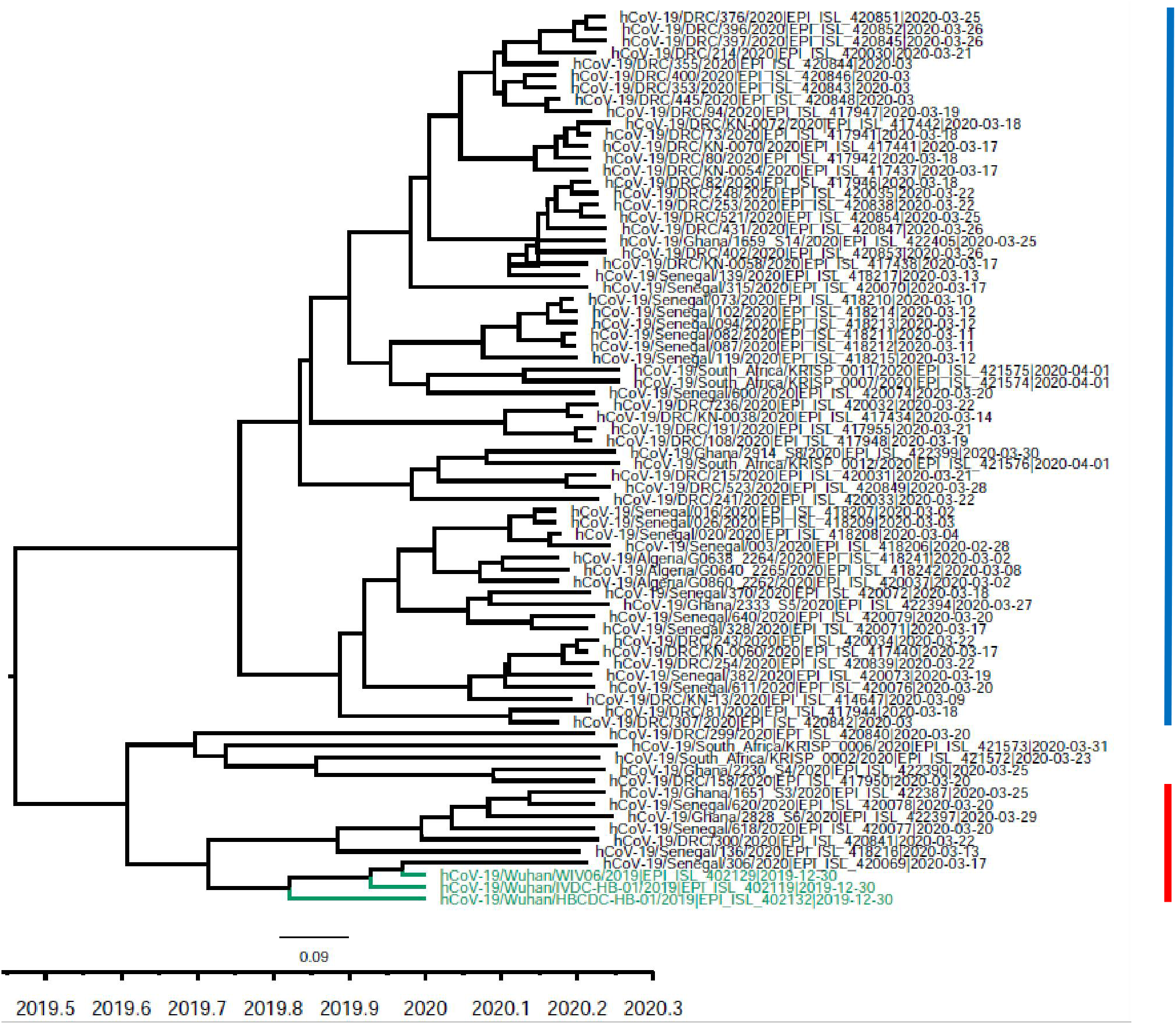
Time scaled MCC tree of AfrSARSCoV-2 sequences, green labels represent reference SARSCoV-2 isolates from Wuhan, China. The blue horizontal line represents lineage B isolates, while the red horizontal line represents lineage A isolates.

**FIGURE 4.**
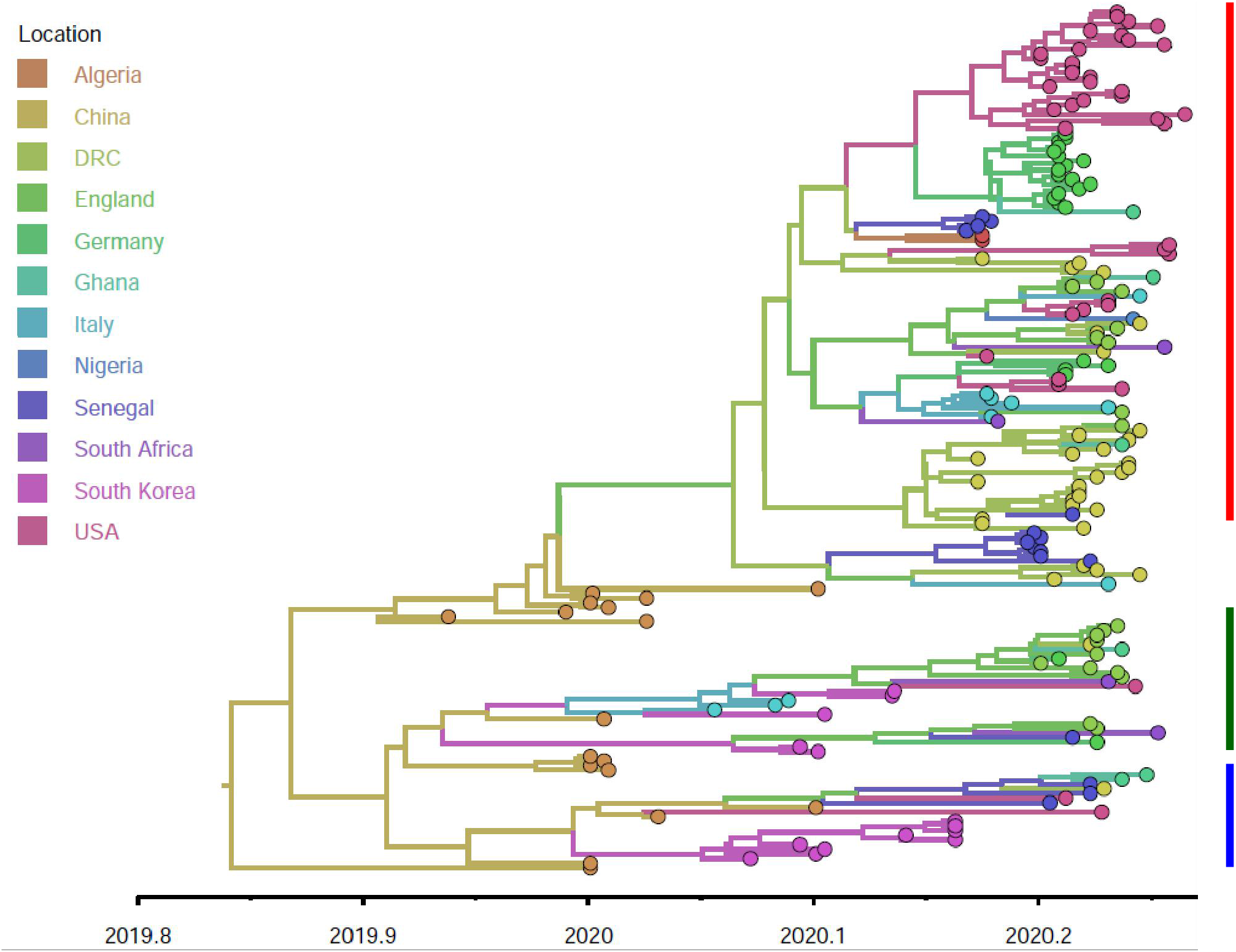
Time scaled MCC tree of complete genome sequences of AfrSARSCoV-2 isolates along with isolates from Asia, Europe and the USA. The legend indicates the color code for each country of origin of the isolates. The red horizontal line represents lineage B.1 isolates, the green horizontal line represents lineage B.2, while the blue horizontal line represents lineage A isolates.

**FIGURE 5.**
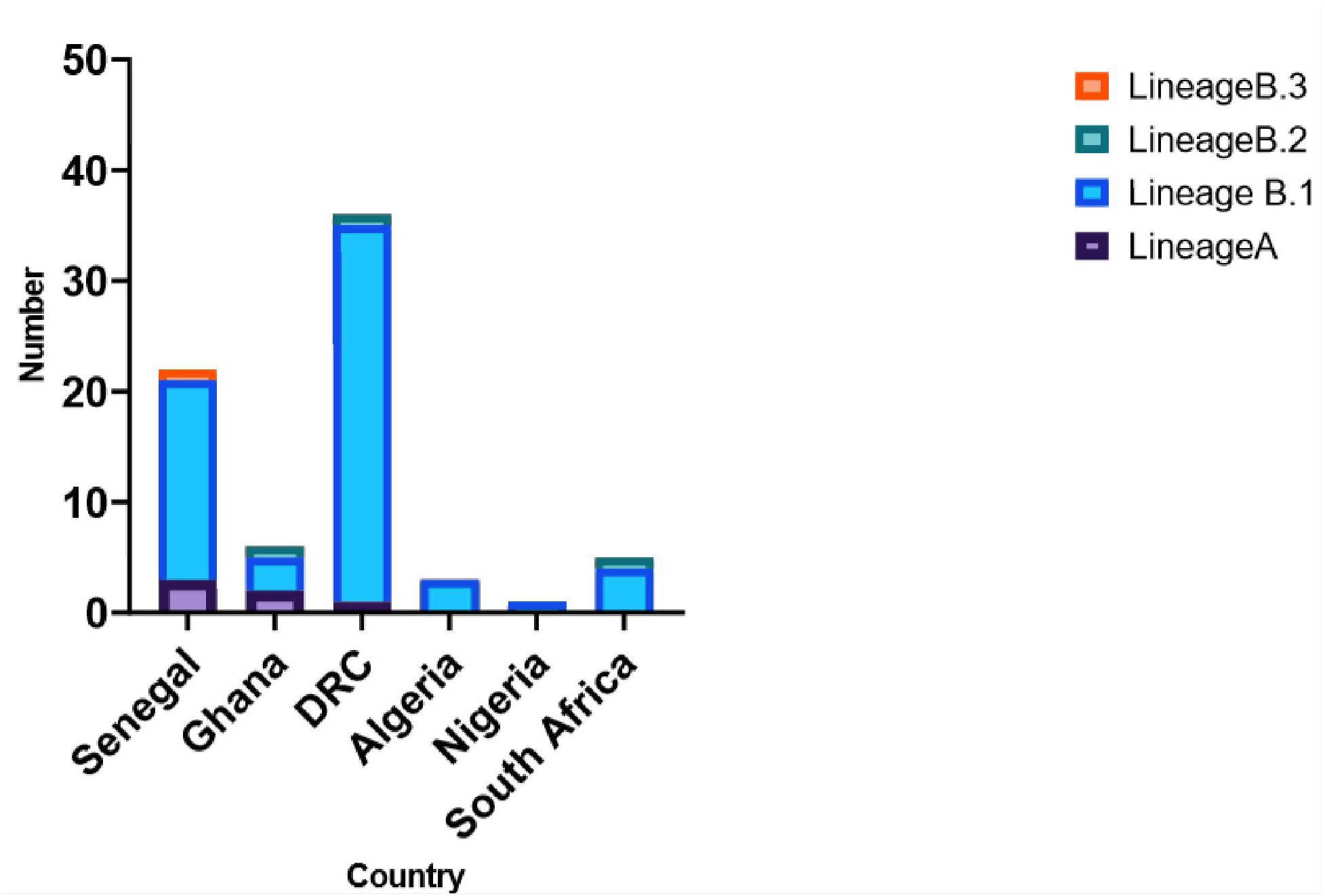
Bar chart showing lineage distribution of AfrSARSCoV-2 sequences according to country of origin.

**FIGURE 6.**
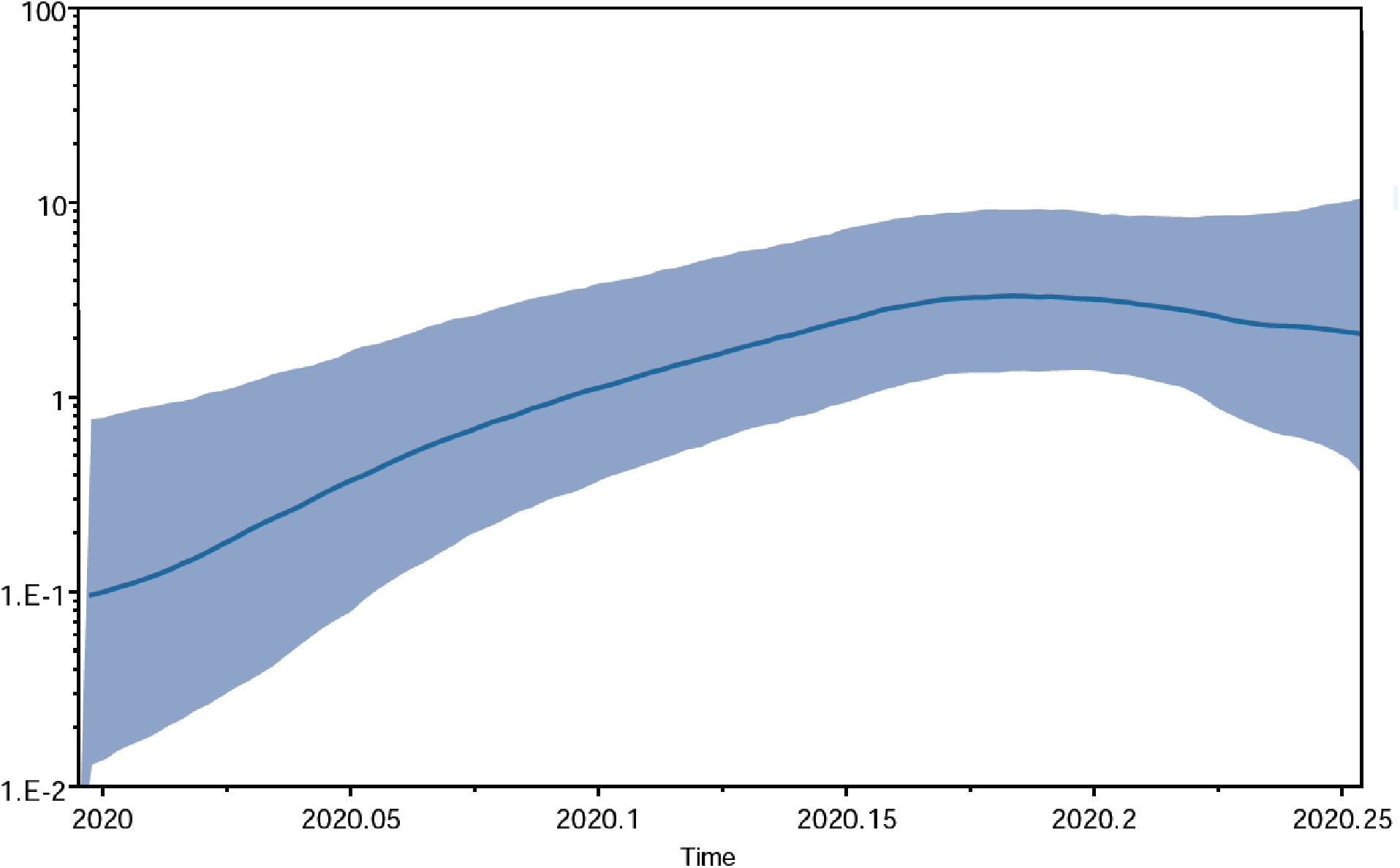
Bayesian Skyline plot of AfrSARSCoV-2 sequences through the early period of the pandemic in Africa.

The AfrSARSCoV-2 sequences were analyzed for the D614-G mutation within the S1 subunit of the spike protein, which has been reported to contribute to increased transmissibility of SARSCoV-2 (Korber et al, 2020). Figure 7 shows a representative amino acid alignment of selected Afr SARSCoV-2 sequences along with reference sequences of BtCoV RaTG13 and PCoV. Our results revealed high prevalence of D614-G mutation among AfrSARSCoV-2 with 12/69 (17.39%). The mutation was recorded in isolates from all African countries analyzed in this study, supplementary figure 2. Prior to this report the D614-G spike mutation was found predominantly in Europe accompanied by high number of cases and significant mortality rate (Pachetti et al, 2020; Korber et al, 2020b). The introduction of this strain in Africa is quite worrisome, considering the population densities of most African cities and the poor state of public health infrastructure to support medical intervention of symptomatic SARSCoV-2 cases. Although more evidence is still required to determine the extent of the effect of the D614-G mutation on the virulence properties of the virus, current evidence from in vitro studies seem to support the hypothesis of increased transmissibility of this variant of the virus (Korber et al, 2020; Hu et al, 2020).

**FIGURE 7.**
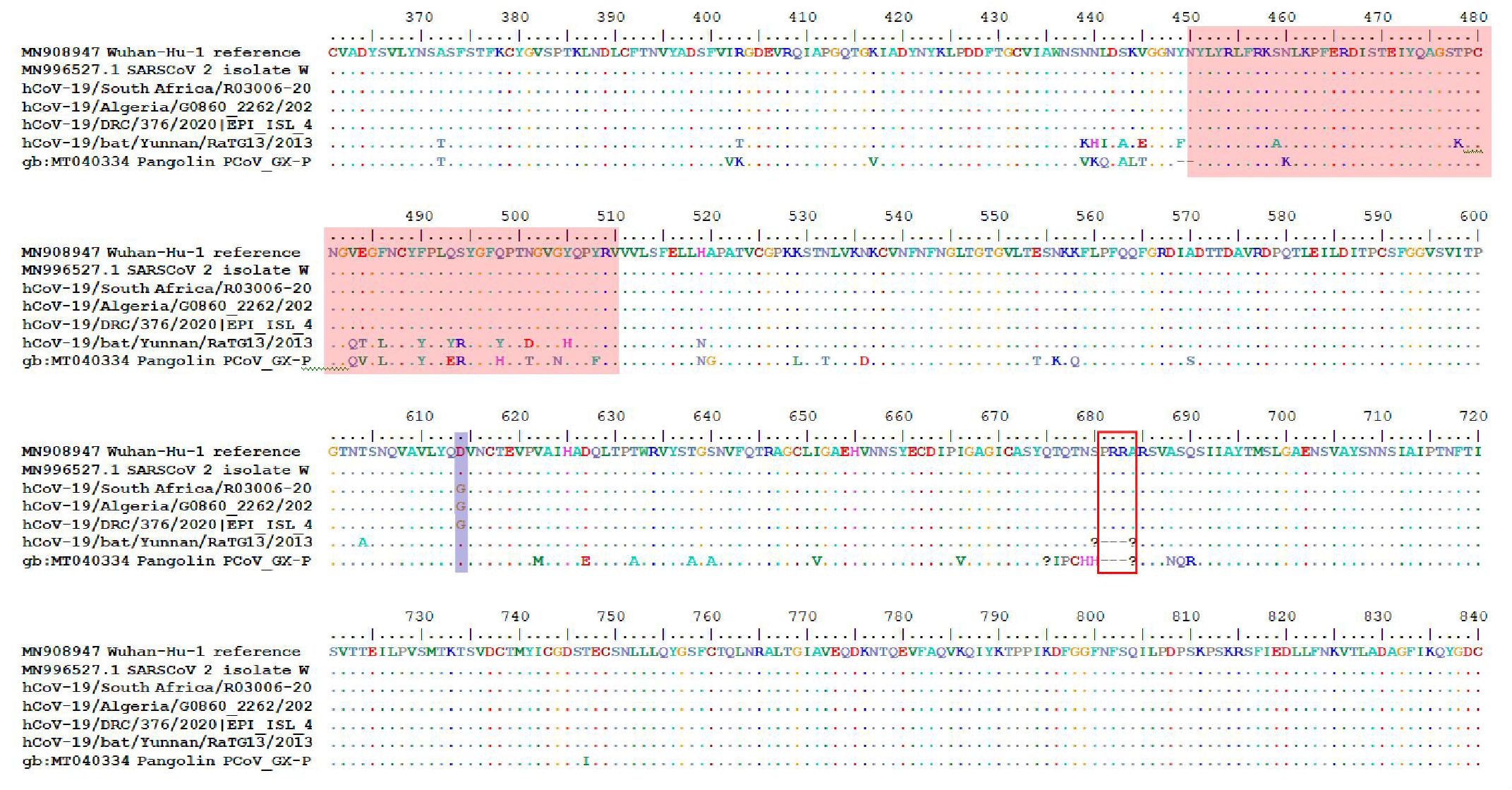
Amino acid alignment of the partial S gene sequences covering amino acid positions 360 to 840, of selected AfrSARSCoV isolates along with reference sequences of closely related PCoV and bat RaTG13. The red shaded region represents the receptor binding domain; the blue shaded box represents the D614-G motive, while the empty red box represents the polybasic cleavage site bordering the S1/S2 sub-unit.

In conclusion we have reported the genetic diversity and evolutionary history of SARSCoV-2 isolated in Africa during the early outbreak period. Our findings have identified diverse sub-lineages of SARSCoV-2 currently circulating among Africans. We also identified high prevalence of the D614-G spike protein variant of the virus capable of rapid transmission in all countries sampled. A major limitation was the relatively low amount of sequence submission available in GISAID database compared with those of other regions such as Europe and Asia. We advocate for upscale of next generation sequencing NGS capacity for whole genome sequencing of SARCoV-2 samples across the African continent to support surveillance and control effort in Africa.

