## Supplementary material for "Evolution and Genetic Diversity of SARSCoV-2 in Africa Using Whole Genome Sequences": Supplimentary table 1

| 1 | hCoV-19/South Africa/R03006-20/2020\|EPI_ISL_417186 |
| --- | --- |
| 2 | hCoV-19/Senegal/600/2020\|EPI_ISL_420074\|2020-03-20 |
| 3 | hCoV-19/Senegal/315/2020\|EPI_ISL_420070\|2020-03-17 |
| 4 | hCoV-19/Senegal/094/2020\|EPI_ISL_418213\|2020-03-12 |
| 5 | hCoV-19/Senegal/087/2020\|EPI_ISL_418212\|2020-03-11 |
| 6 | hCoV-19/Senegal/082/2020\|EPI_ISL_418211\|2020-03-11 |
| 7 | hCoV-19/Senegal/073/2020\|EPI_ISL_418210\|2020-03-10 |
| 8 | hCoV-19/Senegal/003/2020\|EPI_ISL_418206\|2020-02-28 |
| 9 | hCoV-19/Senegal/016/2020\|EPI_ISL_418207\|2020-03-02 |
| 10 | hCoV-19/Senegal/026/2020\|EPI_ISL_418209\|2020-03-03 |
| 11 | hCoV-19/Senegal/020/2020\|EPI_ISL_418208\|2020-03-04 |
| 12 | hCoV-19/Senegal/618/2020\|EPI_ISL_420077\|2020-03-20 |
| 13 | hCoV-19/Senegal/306/2020\|EPI_ISL_420069\|2020-03-17 |
| 14 | hCoV-19/DRC/253/2020\|EPI_ISL_420838\|2020-03-22 |
| 15 | hCoV-19/DRC/431/2020\|EPI_ISL_420847\|2020-03-26 |
| 16 | hCoV-19/DRC/214/2020\|EPI_ISL_420030\|2020-03-21 |
| 17 | hCoV-19/DRC/445/2020\|EPI_ISL_420848\|2020-03 |
| 18 | hCoV-19/DRC/94/2020\|EPI_ISL_417947\|2020-03-19 |
| 19 | hCoV-19/DRC/397/2020\|EPI_ISL_420845\|2020-03-26 |
| 20 | hCoV-19/DRC/396/2020\|EPI_ISL_420852\|2020-03-26 |
| 21 | hCoV-19/DRC/73/2020\|EPI_ISL_417941\|2020-03-18 |
| 22 | hCoV-19/DRC/108/2020\|EPI_ISL_417948\|2020-03-19 |
| 23 | hCoV-19/DRC/353/2020\|EPI_ISL_420843\|2020-03 |
| 24 | hCoV-19/DRC/158/2020\|EPI_ISL_417950\|2020-03-20 |
| 25 | hCoV-19/Senegal/136/2020\|EPI_ISL_418216\|2020-03-13 |
| 26 | hCoV-19/Senegal/620/2020\|EPI_ISL_420078\|2020-03-20 |
| 27 | hCoV-19/Ghana/1651_S3/2020\|EPI_ISL_422387\|2020-03-25 |
| 28 | hCoV-19/DRC/300/2020\|EPI_ISL_420841\|2020-03-22 |
| 29 | hCoV-19/DRC/523/2020\|EPI_ISL_420849\|2020-03-28 |
| 30 | hCoV-19/DRC/215/2020\|EPI_ISL_420031\|2020-03-21 |
| 31 | hCoV-19/DRC/241/2020\|EPI_ISL_420033\|2020-03-22 |
| 32 | hCoV-19/DRC/81/2020\|EPI_ISL_417944\|2020-03-18 |
| 33 | hCoV-19/DRC/243/2020\|EPI_ISL_420034\|2020-03-22 |
| 34 | hCoV-19/DRC/307/2020\|EPI_ISL_420842\|2020-03 |
| 35 | hCoV-19/Senegal/119/2020\|EPI_ISL_418215\|2020-03-12 |
| 36 | hCoV-19/DRC/191/2020\|EPI_ISL_417955\|2020-03-21 |
| 37 | hCoV-19/DRC/376/2020\|EPI_ISL_420851\|2020-03-25 |
| 38 | hCoV-19/Nigeria/Lagos01/2020\|EPI_ISL_413550\|2020-02-27 |
| 39 | hCoV-19/DRC/KN-0060/2020\|EPI_ISL_417440\|2020-03-17 |
| 40 | hCoV-19/South Africa/KRISP_0012/2020\|EPI_ISL_421576\| |
| 41 | hCoV-19/DRC/KN-0038/2020\|EPI_ISL_417434\|2020-03-14 |
| 42 | hCoV-19/South Africa/KRISP_0012/2020\|EPI_ISL_421576 |
| 43 | hCoV-19/Senegal/102/2020\|EPI_ISL_418214\|2020-03-12 |
| 44 | hCoV-19/South Africa/KRISP_0006/2020\|EPI_ISL_421573\|2020-03-31 |
| 45 | hCoV-19/Ghana/2914_S8/2020\|EPI_ISL_422399\|2020-03-30 |
| 46 | hCoV-19/Ghana/2333_S5/2020\|EPI_ISL_422394\|2020-03-27 |
| 47 | hCoV-19/Algeria/G0638_2264/2020\|EPI_ISL_418241\|2020-03-02 |
| 48 | hCoV-19/Algeria/G0860_2262/2020\|EPI_ISL_420037\|2020-03-02 |
| 49 | hCoV-19/Ghana/2230_S4/2020\|EPI_ISL_422390\|2020-03-25 |
| 50 | hCoV-19/Ghana/1659_S14/2020\|EPI_ISL_422405\|2020-03-25 |
| 51 | hCoV-19/Ghana/2828_S6/2020\|EPI_ISL_422397\|2020-03-29 |
| 52 | hCoV-19/South Africa/KRISP_0002/2020\|EPI_ISL_421572 |
| 53 | hCoV-19/Germany/NRW-44/2020\|EPI_ISL_425139\|2020-03-19 |
| 54 | hCoV-19/Germany/NRW-23/2020\|EPI_ISL_419540\|2020-03-16 |
| 55 | hCoV-19/Germany/NRW-26/2020\|EPI_ISL_419543\|2020-03-15 |
| 56 | hCoV-19/Germany/NRW-33/2020\|EPI_ISL_419550\|2020-03-16 |
| 57 | hCoV-19/Germany/NRW-26/2020\|EPI_ISL_419543\|2020-03-15 |
| 58 | hCoV-19/Germany/NRW-22/2020\|EPI_ISL_419539\|2020-03-15 |
| 59 | hCoV-19/Germany/NRW-24/2020\|EPI_ISL_419541\|2020-03-14 |
| 60 | hCoV-19/Germany/NRW-21/2020\|EPI_ISL_419538\|2020-03-14 |
| 61 | hCoV-19/Germany/NRW-42/2020\|EPI_ISL_425126\|2020-03-18 |
| 62 | hCoV-19/Germany/NRW-40/2020\|EPI_ISL_425124\|2020-03-17 |
| 63 | hCoV-19/Germany/NRW-37/2020\|EPI_ISL_425121\|2020-03-16 |
| 64 | hCoV-19/Germany/NRW-42.2/2020\|EPI_ISL_425127\|2020-03-19 |
| 65 | hCoV-19/Germany/NRW-28/2020\|EPI_ISL_419545\|2020-03-15 |
| 66 | hCoV-19/Germany/NRW-38/2020\|EPI_ISL_425122\|2020-03-16 |
| 67 | hCoV-19/Germany/NRW-42.3/2020\|EPI_ISL_425128\|2020-03-20 |
| 68 | hCoV-19/Germany/NRW-25/2020\|EPI_ISL_419542\|2020-03-15 |
| 69 | hCoV-19/Germany/NRW-32/2020\|EPI_ISL_419549\|2020-03-15 |
| 70 | hCoV-19/Germany/NRW-27/2020\|EPI_ISL_419544\|2020-03-15 |
| 71 | hCoV-19/England/SHEF-C0646/2020\|EPI_ISL_420278\|2020-03-21 |
| 72 | hCoV-19/England/SHEF-C06FB/2020\|EPI_ISL_420264\|2020-03-23 |
| 73 | hCoV-19/England/SHEF-C0619/2020\|EPI_ISL_420289\|2020-03-17 |
| 74 | hCoV-19/England/SHEF-C05A3/2020\|EPI_ISL_420279\|2020-03-21 |
| 75 | hCoV-19/England/SHEF-C01EB/2020\|EPI_ISL_420254\|2020-03-25 |
| 76 | hCoV-19/England/SHEF-C0752/2020\|EPI_ISL_420262\|2020-03-24 |
| 77 | hCoV-19/England/SHEF-C06BF/2020\|EPI_ISL_420248\|2020-03-25 |
| 78 | hCoV-19/England/SHEF-C00B1/2020\|EPI_ISL_420276\|2020-03-21 |
| 79 | hCoV-19/England/SHEF-C0066/2020\|EPI_ISL_420277\|2020-03-21 |
| 80 | hCoV-19/England/SHEF-C00EE/2020\|EPI_ISL_420274\|2020-03-22 |
| 81 | hCoV-19/England/SHEF-C04E2/2020\|EPI_ISL_420263\|2020-03-24 |
| 82 | hCoV-19/England/SHEF-C0655/2020\|EPI_ISL_420292\|2020-03-12 |
| 83 | hCoV-19/England/SHEF-C0172/2020\|EPI_ISL_420259\|2020-03-24 |
| 84 | hCoV-19/England/SHEF-C0163/2020\|EPI_ISL_420255\|2020-03-25 |
| 85 | hCoV-19/England/SHEF-C060A/2020\|EPI_ISL_420285\|2020-03-20 |
| 86 | hCoV-19/England/SHEF-C05C1/2020\|EPI_ISL_420282\|2020-03-21 |
| 87 | hCoV-19/Wuhan-Hu-1/2019\|EPI_ISL_402125\|2019-12-31 |
| 88 | hCoV-19/Italy/SPL1/2020\|EPI_ISL_412974\|2020-01-29 |
| 89 | hCoV-19/Wuhan/IPBCAMS-WH-04/2019\|EPI_ISL_403929\|2019-12-30 |
| 90 | hCoV-19/Wuhan/IVDC-HB-01/2019\|EPI_ISL_402119\|2019-12-30 |
| 91 | hCoV-19/Wuhan/WIV04/2019\|EPI_ISL_402124\|2019-12-30 |
| 92 | hCoV-19/Wuhan/IVDC-HB-envF13-21/2020\|EPI_ISL_408515\|2020-01-01 |
| 93 | hCoV-19/Wuhan/IVDC-HB-envF13-20/2020\|EPI_ISL_408514\|2020-01-01 |
| 94 | hCoV-19/Wuhan/IPBCAMS-WH-01/2019\|EPI_ISL_402123\|2019-12-24 |
| 95 | hCoV-19/Korea/KCDC2004/2020\|EPI_ISL_426164\|2020-02-02 |
| 96 | hCoV-19/Korea/KCDC2007/2020\|EPI_ISL_426169\|2020-02-05 |
| 97 | hCoV-19/Wuhan/WHU01/2020\|EPI_ISL_406716\|2020-01-02 |
| 98 | hCoV-19/Wuhan/WHU02/2020\|EPI_ISL_406717\|2020-01-02 |
| 99 | hCoV-19/Shenzhen/HKU-SZ-002/2020\|EPI_ISL_406030\|2020-01-10 |
| 100 | hCoV-19/South Korea/KUMC04/2020\|EPI_ISL_413514\|2020-02-27 |
| 101 | hCoV-19/South Korea/KUMC05/2020\|EPI_ISL_413515\|2020-02-27 |
| 102 | hCoV-19/South Korea/KUMC06/2020\|EPI_ISL_413516\|2020-02-27 |
| 103 | hCoV-19/South Korea/KUMC03/2020\|EPI_ISL_413513\|2020-02-27 |
| 104 | hCoV-19/South Korea/KUMC02/2020\|EPI_ISL_413018\|2020-02-06 |
| 105 | hCoV-19/Korea/KCDC2018/2020\|EPI_ISL_427812\|2020-02-19 |
| 106 | hCoV-19/South Korea/KCDC03/2020\|EPI_ISL_407193\|2020-01-25 |
| 107 | hCoV-19/South Korea/KUMC01/2020\|EPI_ISL_413017\|2020-02-06 |
| 108 | hCoV-19/Korea/KCDC2006/2020\|EPI_ISL_426168\|2020-02-02 |
| 109 | hCoV-19/Korea/KCDC2008/2020\|EPI_ISL_426171\|2020-02-05 |
| 110 | hCoV-19/Wuhan/WIV02/2019\|EPI_ISL_402127\|2019-12-30 |
| 111 | hCoV-19/China/WF0003/2020\|EPI_ISL_413693\|2020-01 |
| 112 | hCoV-19/China/WH-09/2020\|EPI_ISL_411957\|2020-01-08 |
| 113 | hCoV-19/Italy/INMI5/2020\|EPI_ISL_417923\|2020-03-04 |
| 114 | hCoV-19/Italy/INMI1-cs/2020\|EPI_ISL_410546\|2020-01-31 |
| 115 | hCoV-19/Korea/KCDC2016/2020\|EPI_ISL_427810\|2020-02-18 |
| 116 | hCoV-19/Korea/KCDC2017/2020\|EPI_ISL_427811\|2020-02-18 |
| 117 | hCoV-19/Wuhan/WH04/2020\|EPI_ISL_406801\|2020-01-05 |
| 118 | hCoV-19/Wuhan/WH01/2019\|EPI_ISL_406798\|2019-12-26 |
| 119 | hCoV-19/Wuhan/IPBCAMS-WH-02/2019\|EPI_ISL_403931\|2019-12-30 |
| 120 | hCoV-19/England/SHEF-C06DD/2020\|EPI_ISL_420250\|2020-03-25 |
| 121 | hCoV-19/Italy/INMI4/2020\|EPI_ISL_417922\|2020-02-28 |
| 122 | hCoV-19/USA/AZ-ASU2936/2020\|EPI_ISL_424671\|2020-03-17 |
| 123 | hCoV-19/Germany/NRW-48/2020\|EPI_ISL_425132\|2020-03-23 |
| 124 | hCoV-19/USA/AZ-TG269863/2020\|EPI_ISL_426529\|2020-03-16 |
| 125 | hCoV-19/USA/AZ-ASU2922/2020\|EPI_ISL_424668\|2020-03-16 |
| 126 | hCoV-19/Germany/NRW-46/2020\|EPI_ISL_425130\|2020-03-21 |
| 130 | hCoV-19/England/SHEF-C0576/2020\|EPI_ISL_420284\|2020-03-20 |
| 131 | hCoV-19/England/SHEF-C06A0/2020\|EPI_ISL_420280\|2020-03-21 |
| 132 | hCoV-19/USA/AZ_4811/2020\|EPI_ISL_420784\|2020-03-02 |
| 133 | hCoV-19/USA/AZ-TG271856/2020\|EPI_ISL_426538\|2020-03-19 |
| 134 | hCoV-19/USA/AZ-TG268350/2020\|EPI_ISL_426506\|2020-03-17 |
| 135 | hCoV-19/USA/AZ-TG271866/2020\|EPI_ISL_427271\|2020-03-23 |
| 136 | hCoV-19/USA/AZ-TG271868/2020\|EPI_ISL_427272\|2020-03-23 |
| 137 | hCoV-19/USA/AZ-TG271870/2020\|EPI_ISL_426541\|2020-03-24 |
| 138 | hCoV-19/USA/AZ-TG272213/2020\|EPI_ISL_426556\|2020-04-01 |
| 139 | hCoV-19/USA/AZ-TG268915/2020\|EPI_ISL_426518\|2020-03-12 |
| 140 | hCoV-19/USA/AZ-TG269439/2020\|EPI_ISL_426524\|2020-03-20 |
| 141 | hCoV-19/USA/AZ-TG270200/2020\|EPI_ISL_426533\|2020-03-26 |
| 142 | hCoV-19/USA/AZ-TG271591/2020\|EPI_ISL_426536\|2020-03-25 |
| 143 | hCoV-19/USA/AZ-TG269883/2020\|EPI_ISL_426531\|2020-03-12 |
| 144 | hCoV-19/USA/AZ-TG268629/2020\|EPI_ISL_426509\|2020-03-18 |
| 145 | hCoV-19/USA/AZ-TG268911/2020\|EPI_ISL_426516\|2020-03-13 |
| 146 | hCoV-19/USA/AZ-TG273280/2020\|EPI_ISL_426558\|2020-03-25 |
| 147 | hCoV-19/USA/AZ-TG273282/2020\|EPI_ISL_426559\|2020-03-25 |
| 148 | hCoV-19/USA/AZ-TG269440/2020\|EPI_ISL_426525\|2020-03-19 |
| 149 | hCoV-19/USA/AZ-TG268917/2020\|EPI_ISL_426519\|2020-03-14 |
| 150 | hCoV-19/USA/AZ-TG271892/2020\|EPI_ISL_426549\|2020-03-26 |
| 151 | hCoV-19/USA/AZ-TG269027/2020\|EPI_ISL_426520\|2020-03-20 |
| 152 | hCoV-19/USA/AZ-TG269290/2020\|EPI_ISL_426522\|2020-03-20 |
| 153 | hCoV-19/USA/AZ-TG268183/2020\|EPI_ISL_426501\|2020-03-17 |
| 154 | hCoV-19/USA/AZ-TG271578/2020\|EPI_ISL_426535\|2020-03-17 |
| 155 | hCoV-19/USA/AZ-TG268289/2020\|EPI_ISL_426505\|2020-03-17 |
| 156 | hCoV-19/USA/AZ-TG273318/2020\|EPI_ISL_426569\|2020-04-02 |
| 157 | hCoV-19/USA/AZ-TG273304/2020\|EPI_ISL_426564\|2020-04-01 |
| 158 | hCoV-19/USA/AZ-TG273316/2020\|EPI_ISL_426568\|2020-04-02 |
| 159 | hCoV-19/USA/AZ-TG272215/2020\|EPI_ISL_426557\|2020-04-01 |
| 160 | hCoV-19/USA/AZ-TG273300/2020\|EPI_ISL_426562\|2020-03-31 |
| 161 | hCoV-19/USA/AZ-TG273310/2020\|EPI_ISL_426566\|2020-04-01 |
| 162 | hCoV-19/Italy/INMI1-isl/2020\|EPI_ISL_410545\|2020-01-29 |
| 163 | hCoV-19/Germany/NRW-31/2020\|EPI_ISL_419548\|2020-03-15 |
| 164 | hCoV-19/USA/AK-PHL115/2020\|EPI_ISL_427621\|2020-03-25 |
| 165 | hCoV-19/Jingzhou/HBCDC-HB-01/2020\|EPI_ISL_412459\|2020-01-08 |
| 166 | hCoV-19/Italy/INMI10/2020\|EPI_ISL_424344\|2020-03-04 |
| 167 | hCoV-19/Italy/INMI6/2020\|EPI_ISL_419254\|2020-03-23 |
| 168 | hCoV-19/Italy/INMI8/2020\|EPI_ISL_424342\|2020-03-07 |
| 169 | hCoV-19/Italy/INMI9/2020\|EPI_ISL_424343\|2020-03-23 |
| 170 | hCoV-19/Italy/INMI7/2020\|EPI_ISL_419255\|2020-03-23 |
| 171 | hCoV-19/USA/AZ1/2020\|EPI_ISL_406223\|2020-01-22 |
| 172 | hCoV-19/USA/AK-PHL002/2020\|EPI_ISL_420303\|2020-03-15 |
| 173 | hCoV-19/USA/AK-PHL003/2020\|EPI_ISL_420304\|2020-03-15 |
| 174 | hCoV-19/USA/AK-PHL101/2020\|EPI_ISL_427620\|2020-04-04 |
| 175 | hCoV-19/USA/AK-PHL118/2020\|EPI_ISL_427622\|2020-03-24 |
| 176 | hCoV-19/USA/AK-PHL037/2020\|EPI_ISL_424346\|2020-03-21 |
| 177 | hCoV-19/DRC/355/2020\|EPI_ISL_420844\|2020-03 |
| 178 | hCoV-19/DRC/248/2020\|EPI_ISL_420035\|2020-03-22 |
| 179 | hCoV-19/DRC/236/2020\|EPI_ISL_420032\|2020-03-22 |
| 180 | hCoV-19/DRC/400/2020\|EPI_ISL_420846\|2020-03 |
| 181 | hCoV-19/DRC/KN-0072/2020\|EPI_ISL_417442\|2020-03-18 |
| 182 | hCoV-19/DRC/82/2020\|EPI_ISL_417946\|2020-03-18 |
| 183 | hCoV-19/DRC/KN-0070/2020\|EPI_ISL_417441\|2020-03-17 |
| 184 | hCoV-19/DRC/KN-0054/2020\|EPI_ISL_417437\|2020-03-17 |
| 185 | hCoV-19/DRC/80/2020\|EPI_ISL_417942\|2020-03-18 |
| 186 | hCoV-19/DRC/KN-0058/2020\|EPI_ISL_417438\|2020-03-17 |
| 187 | hCoV-19/DRC/352/2020\|EPI_ISL_420850\|2020-03 |
