## Supplementary material for "Evolution and Genetic Diversity of SARSCoV-2 in Africa Using Whole Genome Sequences": Supplimentary table 2

Sequence name	Lineage	Bootstrap	SH-aLRT	Most common countries	Number of taxa	Date range	Days since last sampling

hCoV-19/South	B.1	97	100	"UK, USA, Australia"	6733	"January-24, April-23"	26

hCoV-19/Senegal/600/2020|EPI_ISL_420074|2020-03-20	B.1	98	100	"UK, USA, Australia"	6733	"January-24, April-23"	26

hCoV-19/Senegal/315/2020|EPI_ISL_420070|2020-03-17	B.1	100	100	"UK, USA, Australia"	6733	"January-24, April-23"	26

hCoV-19/Senegal/094/2020|EPI_ISL_418213|2020-03-12	B.1	97	100	"UK, USA, Australia"	6733	"January-24, April-23"	26

hCoV-19/Senegal/087/2020|EPI_ISL_418212|2020-03-11	B.1	99	100	"UK, USA, Australia"	6733	"January-24, April-23"	26

hCoV-19/Senegal/082/2020|EPI_ISL_418211|2020-03-11	B.1	99	100	"UK, USA, Australia"	6733	"January-24, April-23"	26

hCoV-19/Senegal/073/2020|EPI_ISL_418210|2020-03-10	B.1	99	100	"UK, USA, Australia"	6733	"January-24, April-23"	26

hCoV-19/Senegal/003/2020|EPI_ISL_418206|2020-02-28	B.1	100	100	"UK, USA, Australia"	6733	"January-24, April-23"	26

hCoV-19/Senegal/016/2020|EPI_ISL_418207|2020-03-02	B.1	100	100	"UK, USA, Australia"	6733	"January-24, April-23"	26

hCoV-19/Senegal/026/2020|EPI_ISL_418209|2020-03-03	B.1	100	100	"UK, USA, Australia"	6733	"January-24, April-23"	26

hCoV-19/Senegal/020/2020|EPI_ISL_418208|2020-03-04	B.1	100	100	"UK, USA, Australia"	6733	"January-24, April-23"	26

hCoV-19/Senegal/618/2020|EPI_ISL_420077|2020-03-20	A	100	100	"China, South_Korea, USA"	241	"January-05, April-06"	43

hCoV-19/Senegal/306/2020|EPI_ISL_420069|2020-03-17	B.3	97	100	"UK, Denmark, Australia"	469	"February-02, April-19"	30

hCoV-19/DRC/253/2020|EPI_ISL_420838|2020-03-22	B.1	100	100	"UK, USA, Australia"	6733	"January-24, April-23"	26

hCoV-19/DRC/431/2020|EPI_ISL_420847|2020-03-26	B.1	100	100	"UK, USA, Australia"	6733	"January-24, April-23"	26

hCoV-19/DRC/214/2020|EPI_ISL_420030|2020-03-21	B.1.6	99	95	"Belgium, DRC, Australia"	20	"March-11, April-13"	36

hCoV-19/DRC/445/2020|EPI_ISL_420848|2020-03	B.1	100	100	"UK, USA, Australia"	6733	"January-24, April-23"	26

hCoV-19/DRC/94/2020|EPI_ISL_417947|2020-03-19	B.1	100	100	"UK, USA, Australia"	6733	"January-24, April-23"	26

hCoV-19/DRC/397/2020|EPI_ISL_420845|2020-03-26	B.1.6	100	100	"Belgium, DRC, Australia"	20	"March-11, April-13"	36

hCoV-19/DRC/396/2020|EPI_ISL_420852|2020-03-26	B.1.6	100	100	"Belgium, DRC, Australia"	20	"March-11, April-13"	36

hCoV-19/DRC/73/2020|EPI_ISL_417941|2020-03-18	B.1	100	100	"UK, USA, Australia"	6733	"January-24, April-23"	26

hCoV-19/DRC/108/2020|EPI_ISL_417948|2020-03-19	B.1	97	100	"UK, USA, Australia"	6733	"January-24, April-23"	26

hCoV-19/DRC/353/2020|EPI_ISL_420843|2020-03	B.1	100	100	"UK, USA, Australia"	6733	"January-24, April-23"	26

hCoV-19/DRC/158/2020|EPI_ISL_417950|2020-03-20	B.2.1	98	100	"UK, Australia, USA"	919	"February-07, April-22"	27

hCoV-19/Senegal/136/2020|EPI_ISL_418216|2020-03-13	A.2	100	100	"Australia, Spain, UK"	180	"February-26, April-13"	36

hCoV-19/Senegal/620/2020|EPI_ISL_420078|2020-03-20	A	98	100	"China, South_Korea, USA"	241	"January-05, April-06"	43

hCoV-19/Ghana/1651_S3/2020|EPI_ISL_422387|2020-03-25	A	98	94	"China, South_Korea, USA"	241	"January-05, April-06"	43

hCoV-19/DRC/300/2020|EPI_ISL_420841|2020-03-22	A	98	100	"China, South_Korea, USA"	241	"January-05, April-06"	43

hCoV-19/DRC/523/2020|EPI_ISL_420849|2020-03-28	B.1.1	100	100	"UK, Iceland, Russia"	329	"February-07, April-22"	27

hCoV-19/DRC/215/2020|EPI_ISL_420031|2020-03-21	B.1.1	100	100	"UK, Iceland, Russia"	329	"February-07, April-22"	27

hCoV-19/DRC/241/2020|EPI_ISL_420033|2020-03-22	B.1	100	100	"UK, USA, Australia"	6733	"January-24, April-23"	26

hCoV-19/DRC/81/2020|EPI_ISL_417944|2020-03-18	B.1	100	100	"UK, USA, Australia"	6733	"January-24, April-23"	26

hCoV-19/DRC/243/2020|EPI_ISL_420034|2020-03-22	B.1	100	100	"UK, USA, Australia"	6733	"January-24, April-23"	26

hCoV-19/DRC/307/2020|EPI_ISL_420842|2020-03	B.1	100	100	"UK, USA, Australia"	6733	"January-24, April-23"	26

hCoV-19/Senegal/119/2020|EPI_ISL_418215|2020-03-12	B.1	98	100	"UK, USA, Australia"	6733	"January-24, April-23"	26

hCoV-19/DRC/191/2020|EPI_ISL_417955|2020-03-21	B.1	96	95	"UK, USA, Australia"	6733	"January-24, April-23"	26

hCoV-19/DRC/376/2020|EPI_ISL_420851|2020-03-25	B.1.6	100	100	"Belgium, DRC, Australia"	20	"March-11, April-13"	36

hCoV-19/Nigeria/Lagos01/2020|EPI_ISL_413550|2020-02-27	B.1	100	100	"UK, USA, Australia"	6733	"January-24, April-23"	26

hCoV-19/DRC/KN-0060/2020|EPI_ISL_417440|2020-03-17	B.1	100	100	"UK, USA, Australia"	6733	"January-24, April-23"	26

hCoV-19/DRC/KN-0038/2020|EPI_ISL_417434|2020-03-14	B.1	99	100	"UK, USA, Australia"	6733	"January-24, April-23"	26

hCoV-19/South	B.1.10	91	91	"UK, Iceland, Australia"	65	"March-06, April-18"	31

hCoV-19/Senegal/102/2020|EPI_ISL_418214|2020-03-12	B.1	98	100	"UK, USA, Australia"	6733	"January-24, April-23"	26

hCoV-19/South	B.6	100	96	"Australia, Singapore, USA"	94	"March-04, April-15"	34

hCoV-19/Ghana/2914_S8/2020|EPI_ISL_422399|2020-03-30	B.1	100	100	"UK, USA, Australia"	6733	"January-24, April-23"	26

hCoV-19/Ghana/2333_S5/2020|EPI_ISL_422394|2020-03-27	B.1.3	100	100	"USA, Australia, Argentina"	173	"March-10, April-18"	31

hCoV-19/Algeria/G0638_2264/2020|EPI_ISL_418241|2020-03-02	B.1	100	100	"UK, USA, Australia"	6733	"January-24, April-23"	26

hCoV-19/Algeria/G0860_2262/2020|EPI_ISL_420037|2020-03-02	B.1	100	100	"UK, USA, Australia"	6733	"January-24, April-23"	26

hCoV-19/Ghana/2230_S4/2020|EPI_ISL_422390|2020-03-25	B.2.1	96	100	"UK, Australia, USA"	919	"February-07, April-22"	27

hCoV-19/Ghana/1659_S14/2020|EPI_ISL_422405|2020-03-25	B.1	100	100	"UK, USA, Australia"	6733	"January-24, April-23"	26

hCoV-19/Ghana/2828_S6/2020|EPI_ISL_422397|2020-03-29	A	100	100	"China, South_Korea, USA"	241	"January-05, April-06"	43

hCoV-19/South	B.2.2	88	100	"UK, Australia, Iceland"	128	"February-25, April-13"	36

hCoV-19/Germany/NRW-44/2020|EPI_ISL_425139|2020-03-19	B.1	95	100	"UK, USA, Australia"	6733	"January-24, April-23"	26

hCoV-19/Germany/NRW-23/2020|EPI_ISL_419540|2020-03-16	B.1	100	100	"UK, USA, Australia"	6733	"January-24, April-23"	26

hCoV-19/Germany/NRW-26/2020|EPI_ISL_419543|2020-03-15	B.1	100	100	"UK, USA, Australia"	6733	"January-24, April-23"	26

hCoV-19/Germany/NRW-34/2020|EPI_ISL_419551|2020-03-16	B.1.8	100	100	"Netherlands, Iceland, Australia"	76	"March-04, April-03"	46

hCoV-19/Germany/NRW-33/2020|EPI_ISL_419550|2020-03-16	B.1.8	100	100	"Netherlands, Iceland, Australia"	76	"March-04, April-03"	46

hCoV-19/Germany/NRW-23/2020|EPI_ISL_419540|2020-03-16(2)	B.1	100	100	"UK, USA, Australia"	6733	"January-24, April-23"	26

hCoV-19/Germany/NRW-26/2020|EPI_ISL_419543|2020-03-15(2)	B.1	100	100	"UK, USA, Australia"	6733	"January-24, April-23"	26

hCoV-19/Germany/NRW-22/2020|EPI_ISL_419539|2020-03-15	B.1	100	100	"UK, USA, Australia"	6733	"January-24, April-23"	26

hCoV-19/Germany/NRW-24/2020|EPI_ISL_419541|2020-03-14	B.1	100	100	"UK, USA, Australia"	6733	"January-24, April-23"	26

hCoV-19/Germany/NRW-21/2020|EPI_ISL_419538|2020-03-14	B.1	100	100	"UK, USA, Australia"	6733	"January-24, April-23"	26

hCoV-19/Germany/NRW-42/2020|EPI_ISL_425126|2020-03-18	B.1	100	100	"UK, USA, Australia"	6733	"January-24, April-23"	26

hCoV-19/Germany/NRW-40/2020|EPI_ISL_425124|2020-03-17	B.1	100	100	"UK, USA, Australia"	6733	"January-24, April-23"	26

hCoV-19/Germany/NRW-37/2020|EPI_ISL_425121|2020-03-16	B.1	100	100	"UK, USA, Australia"	6733	"January-24, April-23"	26

hCoV-19/Germany/NRW-42.2/2020|EPI_ISL_425127|2020-03-19	B.1	100	100	"UK, USA, Australia"	6733	"January-24, April-23"	26

hCoV-19/Germany/NRW-28/2020|EPI_ISL_419545|2020-03-15	B.1	100	100	"UK, USA, Australia"	6733	"January-24, April-23"	26

hCoV-19/Germany/NRW-38/2020|EPI_ISL_425122|2020-03-16	B.1	100	100	"UK, USA, Australia"	6733	"January-24, April-23"	26

hCoV-19/Germany/NRW-42.3/2020|EPI_ISL_425128|2020-03-20	B.1	100	100	"UK, USA, Australia"	6733	"January-24, April-23"	26

hCoV-19/Germany/NRW-25/2020|EPI_ISL_419542|2020-03-15	B.1	100	100	"UK, USA, Australia"	6733	"January-24, April-23"	26

hCoV-19/Germany/NRW-32/2020|EPI_ISL_419549|2020-03-15	B.1	100	100	"UK, USA, Australia"	6733	"January-24, April-23"	26

hCoV-19/Germany/NRW-27/2020|EPI_ISL_419544|2020-03-15	B.1	100	100	"UK, USA, Australia"	6733	"January-24, April-23"	26

hCoV-19/England/SHEF-C0646/2020|EPI_ISL_420278|2020-03-21	B.1.1	100	100	"UK, Iceland, Russia"	329	"February-07, April-22"	27

hCoV-19/England/SHEF-C06FB/2020|EPI_ISL_420264|2020-03-23	B.1.1	100	100	"UK, Iceland, Russia"	329	"February-07, April-22"	27

hCoV-19/England/SHEF-C0619/2020|EPI_ISL_420289|2020-03-17	B.1	100	100	"UK, USA, Australia"	6733	"January-24, April-23"	26

hCoV-19/England/SHEF-C05A3/2020|EPI_ISL_420279|2020-03-21	B.1	100	100	"UK, USA, Australia"	6733	"January-24, April-23"	26

hCoV-19/England/SHEF-C01EB/2020|EPI_ISL_420254|2020-03-25	B.1	100	100	"UK, USA, Australia"	6733	"January-24, April-23"	26

hCoV-19/England/SHEF-C0752/2020|EPI_ISL_420262|2020-03-24	B.1.1	100	100	"UK, Iceland, Russia"	329	"February-07, April-22"	27

hCoV-19/England/SHEF-C06BF/2020|EPI_ISL_420248|2020-03-25	B.1.5	85	100	"UK, Spain, Australia"	424	"February-26, April-23"	26

hCoV-19/England/SHEF-C00B1/2020|EPI_ISL_420276|2020-03-21	B.2.1	98	100	"UK, Australia, USA"	919	"February-07, April-22"	27

hCoV-19/England/SHEF-C0066/2020|EPI_ISL_420277|2020-03-21	B.2.1	95	100	"UK, Australia, USA"	919	"February-07, April-22"	27

hCoV-19/England/SHEF-C00EE/2020|EPI_ISL_420274|2020-03-22	B.2.1	98	100	"UK, Australia, USA"	919	"February-07, April-22"	27

hCoV-19/England/SHEF-C04E2/2020|EPI_ISL_420263|2020-03-24	B.2.1	95	100	"UK, Australia, USA"	919	"February-07, April-22"	27

hCoV-19/England/SHEF-C0655/2020|EPI_ISL_420292|2020-03-12	B.2.1	98	100	"UK, Australia, USA"	919	"February-07, April-22"	27

hCoV-19/England/SHEF-C0172/2020|EPI_ISL_420259|2020-03-24	B.2	79	100	"UK, Iceland, USA"	565	"February-13, April-23"	26

hCoV-19/England/SHEF-C0163/2020|EPI_ISL_420255|2020-03-25	B.2	84	100	"UK, Iceland, USA"	565	"February-13, April-23"	26

hCoV-19/England/SHEF-C060A/2020|EPI_ISL_420285|2020-03-20	B.2.1	82	100	"UK, Australia, USA"	919	"February-07, April-22"	27

hCoV-19/England/SHEF-C05C1/2020|EPI_ISL_420282|2020-03-21	B.2.1	98	100	"UK, Australia, USA"	919	"February-07, April-22"	27

hCoV-19/Wuhan-Hu-1/2019|EPI_ISL_402125|2019-12-31	B	99	100	"UK, China, USA"	1016	"December-24, April-23"	26

hCoV-19/Italy/SPL1/2020|EPI_ISL_412974|2020-01-29	B	100	100	"UK, China, USA"	1016	"December-24, April-23"	26

hCoV-19/Wuhan/IPBCAMS-WH-04/2019|EPI_ISL_403929|2019-12-30	B	99	100	"UK, China, USA"	1016	"December-24, April-23"	26

hCoV-19/Wuhan/IVDC-HB-01/2019|EPI_ISL_402119|2019-12-30	B	99	100	"UK, China, USA"	1016	"December-24, April-23"	26

hCoV-19/Wuhan/WIV04/2019|EPI_ISL_402124|2019-12-30	B	99	100	"UK, China, USA"	1016	"December-24, April-23"	26

hCoV-19/Wuhan/IVDC-HB-envF13-21/2020|EPI_ISL_408515|2020-01-01	B	97	93	"UK, China, USA"	1016	"December-24, April-23"	26

hCoV-19/Wuhan/IVDC-HB-envF13-20/2020|EPI_ISL_408514|2020-01-01	B	99	100	"UK, China, USA"	1016	"December-24, April-23"	26

hCoV-19/Wuhan/IPBCAMS-WH-01/2019|EPI_ISL_402123|2019-12-24	B	94	90	"UK, China, USA"	1016	"December-24, April-23"	26

hCoV-19/Korea/KCDC2004/2020|EPI_ISL_426164|2020-02-02	B	99	100	"UK, China, USA"	1016	"December-24, April-23"	26

hCoV-19/Korea/KCDC2007/2020|EPI_ISL_426169|2020-02-05	B	99	100	"UK, China, USA"	1016	"December-24, April-23"	26

hCoV-19/Wuhan/WHU01/2020|EPI_ISL_406716|2020-01-02	B	99	100	"UK, China, USA"	1016	"December-24, April-23"	26

hCoV-19/Wuhan/WHU02/2020|EPI_ISL_406717|2020-01-02	B	99	100	"UK, China, USA"	1016	"December-24, April-23"	26

hCoV-19/Shenzhen/HKU-SZ-002/2020|EPI_ISL_406030|2020-01-10	A	100	100	"China, South_Korea, USA"	241	"January-05, April-06"	43

hCoV-19/South	A	100	100	"China, South_Korea, USA"	241	"January-05, April-06"	43

hCoV-19/South	A	100	100	"China, South_Korea, USA"	241	"January-05, April-06"	43

hCoV-19/South	A	100	100	"China, South_Korea, USA"	241	"January-05, April-06"	43

hCoV-19/South	A	100	100	"China, South_Korea, USA"	241	"January-05, April-06"	43

hCoV-19/South	B	98	91	"UK, China, USA"	1016	"December-24, April-23"	26

hCoV-19/Korea/KCDC2018/2020|EPI_ISL_427812|2020-02-19	A	100	100	"China, South_Korea, USA"	241	"January-05, April-06"	43

hCoV-19/South	A	100	100	"China, South_Korea, USA"	241	"January-05, April-06"	43

hCoV-19/South	A	100	100	"China, South_Korea, USA"	241	"January-05, April-06"	43

hCoV-19/Korea/KCDC2006/2020|EPI_ISL_426168|2020-02-02	A	100	100	"China, South_Korea, USA"	241	"January-05, April-06"	43

hCoV-19/Korea/KCDC2008/2020|EPI_ISL_426171|2020-02-05	A	100	100	"China, South_Korea, USA"	241	"January-05, April-06"	43

hCoV-19/Wuhan/WIV02/2019|EPI_ISL_402127|2019-12-30	B	89	92	"UK, China, USA"	1016	"December-24, April-23"	26

hCoV-19/China/WF0003/2020|EPI_ISL_413693|2020-01	B	89	92	"UK, China, USA"	1016	"December-24, April-23"	26

hCoV-19/China/WH-09/2020|EPI_ISL_411957|2020-01-08	B	99	100	"UK, China, USA"	1016	"December-24, April-23"	26

hCoV-19/Italy/INMI5/2020|EPI_ISL_417923|2020-03-04	B.1.5	93	100	"UK, Spain, Australia"	424	"February-26, April-23"	26

hCoV-19/Italy/INMI1-cs/2020|EPI_ISL_410546|2020-01-31	B	100	100	"UK, China, USA"	1016	"December-24, April-23"	26

hCoV-19/Korea/KCDC2016/2020|EPI_ISL_427810|2020-02-18	B.2	92	100	"UK, Iceland, USA"	565	"February-13, April-23"	26

hCoV-19/Korea/KCDC2017/2020|EPI_ISL_427811|2020-02-18	B.2	92	100	"UK, Iceland, USA"	565	"February-13, April-23"	26

hCoV-19/Wuhan/WH04/2020|EPI_ISL_406801|2020-01-05	A	100	100	"China, South_Korea, USA"	241	"January-05, April-06"	43

hCoV-19/Wuhan/WH01/2019|EPI_ISL_406798|2019-12-26	B	89	92	"UK, China, USA"	1016	"December-24, April-23"	26

hCoV-19/Wuhan/IPBCAMS-WH-02/2019|EPI_ISL_403931|2019-12-30	B	99	100	"UK, China, USA"	1016	"December-24, April-23"	26

hCoV-19/England/SHEF-C06DD/2020|EPI_ISL_420250|2020-03-25	B.1	100	100	"UK, USA, Australia"	6733	"January-24, April-23"	26

hCoV-19/Italy/INMI4/2020|EPI_ISL_417922|2020-02-28	B.1	100	100	"UK, USA, Australia"	6733	"January-24, April-23"	26

hCoV-19/USA/AZ-ASU2936/2020|EPI_ISL_424671|2020-03-17	B.1	100	100	"UK, USA, Australia"	6733	"January-24, April-23"	26

hCoV-19/Germany/NRW-48/2020|EPI_ISL_425132|2020-03-23	B.1	98	100	"UK, USA, Australia"	6733	"January-24, April-23"	26

hCoV-19/USA/AZ-TG269863/2020|EPI_ISL_426529|2020-03-16	B.1	100	100	"UK, USA, Australia"	6733	"January-24, April-23"	26

hCoV-19/USA/AZ-ASU2922/2020|EPI_ISL_424668|2020-03-16	A.1	95	100	"USA, Australia, Canada"	981	"February-22, April-09"	40

hCoV-19/Germany/NRW-46/2020|EPI_ISL_425130|2020-03-21	B.4	93	100	"Australia, UK, Canada"	206	"January-18, April-14"	35

hCoV-19/England/SHEF-C0576/2020|EPI_ISL_420284|2020-03-20	B	96	100	"UK, China, USA"	1016	"December-24, April-23"	26

hCoV-19/England/SHEF-C06A0/2020|EPI_ISL_420280|2020-03-21	B	94	100	"UK, China, USA"	1016	"December-24, April-23"	26

hCoV-19/USA/AZ_4811/2020|EPI_ISL_420784|2020-03-02	B.1	100	100	"UK, USA, Australia"	6733	"January-24, April-23"	26

hCoV-19/USA/AZ-TG271856/2020|EPI_ISL_426538|2020-03-19	B.1	100	100	"UK, USA, Australia"	6733	"January-24, April-23"	26

hCoV-19/USA/AZ-TG268350/2020|EPI_ISL_426506|2020-03-17	B.1	100	100	"UK, USA, Australia"	6733	"January-24, April-23"	26

hCoV-19/USA/AZ-TG271870/2020|EPI_ISL_426541|2020-03-24	B.1.2	76	100	"USA, Australia, Canada"	145	"March-09, April-14"	35

hCoV-19/USA/AZ-TG272213/2020|EPI_ISL_426556|2020-04-01	B.1.2	79	100	"USA, Australia, Canada"	145	"March-09, April-14"	35

hCoV-19/USA/AZ-TG268915/2020|EPI_ISL_426518|2020-03-12	B.1.2	79	100	"USA, Australia, Canada"	145	"March-09, April-14"	35

hCoV-19/USA/AZ-TG269439/2020|EPI_ISL_426524|2020-03-20	B.1.2	79	100	"USA, Australia, Canada"	145	"March-09, April-14"	35

hCoV-19/USA/AZ-TG270200/2020|EPI_ISL_426533|2020-03-26	B.1.2	79	100	"USA, Australia, Canada"	145	"March-09, April-14"	35

hCoV-19/USA/AZ-TG271591/2020|EPI_ISL_426536|2020-03-25	B.1.2	74	85	"USA, Australia, Canada"	145	"March-09, April-14"	35

hCoV-19/USA/AZ-TG269883/2020|EPI_ISL_426531|2020-03-12	B.1.2	79	100	"USA, Australia, Canada"	145	"March-09, April-14"	35

hCoV-19/USA/AZ-TG268629/2020|EPI_ISL_426509|2020-03-18	B.1.2	75	85	"USA, Australia, Canada"	145	"March-09, April-14"	35

hCoV-19/USA/AZ-TG268911/2020|EPI_ISL_426516|2020-03-13	B.1.2	75	100	"USA, Australia, Canada"	145	"March-09, April-14"	35

hCoV-19/USA/AZ-TG273280/2020|EPI_ISL_426558|2020-03-25	B.1	100	100	"UK, USA, Australia"	6733	"January-24, April-23"	26

hCoV-19/USA/AZ-TG273282/2020|EPI_ISL_426559|2020-03-25	B.1	100	100	"UK, USA, Australia"	6733	"January-24, April-23"	26

hCoV-19/USA/AZ-TG269440/2020|EPI_ISL_426525|2020-03-19	B.1	100	100	"UK, USA, Australia"	6733	"January-24, April-23"	26

hCoV-19/USA/AZ-TG268917/2020|EPI_ISL_426519|2020-03-14	B.1	100	100	"UK, USA, Australia"	6733	"January-24, April-23"	26

hCoV-19/USA/AZ-TG271892/2020|EPI_ISL_426549|2020-03-26	B.1.2	80	100	"USA, Australia, Canada"	145	"March-09, April-14"	35

hCoV-19/USA/AZ-TG269027/2020|EPI_ISL_426520|2020-03-20	B.1.2	83	100	"USA, Australia, Canada"	145	"March-09, April-14"	35

hCoV-19/USA/AZ-TG269290/2020|EPI_ISL_426522|2020-03-20	B.1.2	83	100	"USA, Australia, Canada"	145	"March-09, April-14"	35

hCoV-19/USA/AZ-TG268183/2020|EPI_ISL_426501|2020-03-17	B.1.2	81	100	"USA, Australia, Canada"	145	"March-09, April-14"	35

hCoV-19/USA/AZ-TG271578/2020|EPI_ISL_426535|2020-03-17	B.1.2	80	100	"USA, Australia, Canada"	145	"March-09, April-14"	35

hCoV-19/USA/AZ-TG268289/2020|EPI_ISL_426505|2020-03-17	B.1.2	80	100	"USA, Australia, Canada"	145	"March-09, April-14"	35

hCoV-19/USA/AZ-TG273318/2020|EPI_ISL_426569|2020-04-02	B.1	100	100	"UK, USA, Australia"	6733	"January-24, April-23"	26

hCoV-19/USA/AZ-TG273304/2020|EPI_ISL_426564|2020-04-01	B.1	100	100	"UK, USA, Australia"	6733	"January-24, April-23"	26

hCoV-19/USA/AZ-TG273316/2020|EPI_ISL_426568|2020-04-02	B.1	100	100	"UK, USA, Australia"	6733	"January-24, April-23"	26

hCoV-19/USA/AZ-TG272215/2020|EPI_ISL_426557|2020-04-01	B.1.2	80	100	"USA, Australia, Canada"	145	"March-09, April-14"	35

hCoV-19/USA/AZ-TG273300/2020|EPI_ISL_426562|2020-03-31	B.1.29	95	100	"USA, Australia"	19	"March-15, April-16"	33

hCoV-19/USA/AZ-TG273310/2020|EPI_ISL_426566|2020-04-01	B.1.29	98	100	"USA, Australia"	19	"March-15, April-16"	33

hCoV-19/Italy/INMI1-isl/2020|EPI_ISL_410545|2020-01-29	B	100	100	"UK, China, USA"	1016	"December-24, April-23"	26

hCoV-19/Germany/NRW-31/2020|EPI_ISL_419548|2020-03-15	B.2	36	0	"UK, Iceland, USA"	565	"February-13, April-23"	26

hCoV-19/USA/AK-PHL115/2020|EPI_ISL_427621|2020-03-25	B.1	98	100	"UK, USA, Australia"	6733	"January-24, April-23"	26

hCoV-19/Jingzhou/HBCDC-HB-01/2020|EPI_ISL_412459|2020-01-08	B	96	92	"UK, China, USA"	1016	"December-24, April-23"	26

hCoV-19/Italy/INMI10/2020|EPI_ISL_424344|2020-03-04	B.1	100	100	"UK, USA, Australia"	6733	"January-24, April-23"	26

hCoV-19/Italy/INMI6/2020|EPI_ISL_419254|2020-03-23	B.1	100	100	"UK, USA, Australia"	6733	"January-24, April-23"	26

hCoV-19/Italy/INMI8/2020|EPI_ISL_424342|2020-03-07	B.1	98	100	"UK, USA, Australia"	6733	"January-24, April-23"	26

hCoV-19/Italy/INMI9/2020|EPI_ISL_424343|2020-03-23	B.1	98	100	"UK, USA, Australia"	6733	"January-24, April-23"	26

hCoV-19/Italy/INMI7/2020|EPI_ISL_419255|2020-03-23	B.1	99	100	"UK, USA, Australia"	6733	"January-24, April-23"	26

hCoV-19/USA/AZ1/2020|EPI_ISL_406223|2020-01-22	A	81	100	"China, South_Korea, USA"	241	"January-05, April-06"	43

hCoV-19/USA/AK-PHL002/2020|EPI_ISL_420303|2020-03-15	B.1	98	100	"UK, USA, Australia"	6733	"January-24, April-23"	26

hCoV-19/USA/AK-PHL003/2020|EPI_ISL_420304|2020-03-15	B.1	98	100	"UK, USA, Australia"	6733	"January-24, April-23"	26

hCoV-19/USA/AK-PHL101/2020|EPI_ISL_427620|2020-04-04	B.1.38	88	91	USA	12	"March-17, April-08"	41

hCoV-19/USA/AK-PHL118/2020|EPI_ISL_427622|2020-03-24	B.1.2	98	100	"USA, Australia, Canada"	145	"March-09, April-14"	35

hCoV-19/USA/AK-PHL037/2020|EPI_ISL_424346|2020-03-21	B.2	100	100	"UK, Iceland, USA"	565	"February-13, April-23"	26

hCoV-19/DRC/355/2020|EPI_ISL_420844|2020-03	B.1	100	100	"UK, USA, Australia"	6733	"January-24, April-23"	26

hCoV-19/DRC/248/2020|EPI_ISL_420035|2020-03-22	B.1	100	100	"UK, USA, Australia"	6733	"January-24, April-23"	26

hCoV-19/DRC/236/2020|EPI_ISL_420032|2020-03-22	B.1	95	95	"UK, USA, Australia"	6733	"January-24, April-23"	26

hCoV-19/DRC/400/2020|EPI_ISL_420846|2020-03	B.1	100	100	"UK, USA, Australia"	6733	"January-24, April-23"	26

hCoV-19/DRC/KN-0072/2020|EPI_ISL_417442|2020-03-18	B.1	100	100	"UK, USA, Australia"	6733	"January-24, April-23"	26

hCoV-19/DRC/82/2020|EPI_ISL_417946|2020-03-18	B.1	100	100	"UK, USA, Australia"	6733	"January-24, April-23"	26

hCoV-19/DRC/KN-0070/2020|EPI_ISL_417441|2020-03-17	B.1	100	100	"UK, USA, Australia"	6733	"January-24, April-23"	26

hCoV-19/DRC/KN-0054/2020|EPI_ISL_417437|2020-03-17	B.1	100	100	"UK, USA, Australia"	6733	"January-24, April-23"	26

hCoV-19/DRC/80/2020|EPI_ISL_417942|2020-03-18	B.1	100	100	"UK, USA, Australia"	6733	"January-24, April-23"	26

hCoV-19/DRC/KN-0058/2020|EPI_ISL_417438|2020-03-17	B.1	100	100	"UK, USA, Australia"	6733	"January-24, April-23"	26

hCoV-19/DRC/352/2020|EPI_ISL_420850|2020-03	B.1	100	100	"UK, USA, Australia"	6733	"January-24, April-23"	26
