## Supplementary figures and images for "Evolution and Genetic Diversity of SARSCoV-2 in Africa Using Whole Genome Sequences"

### Supplimentary figure 1

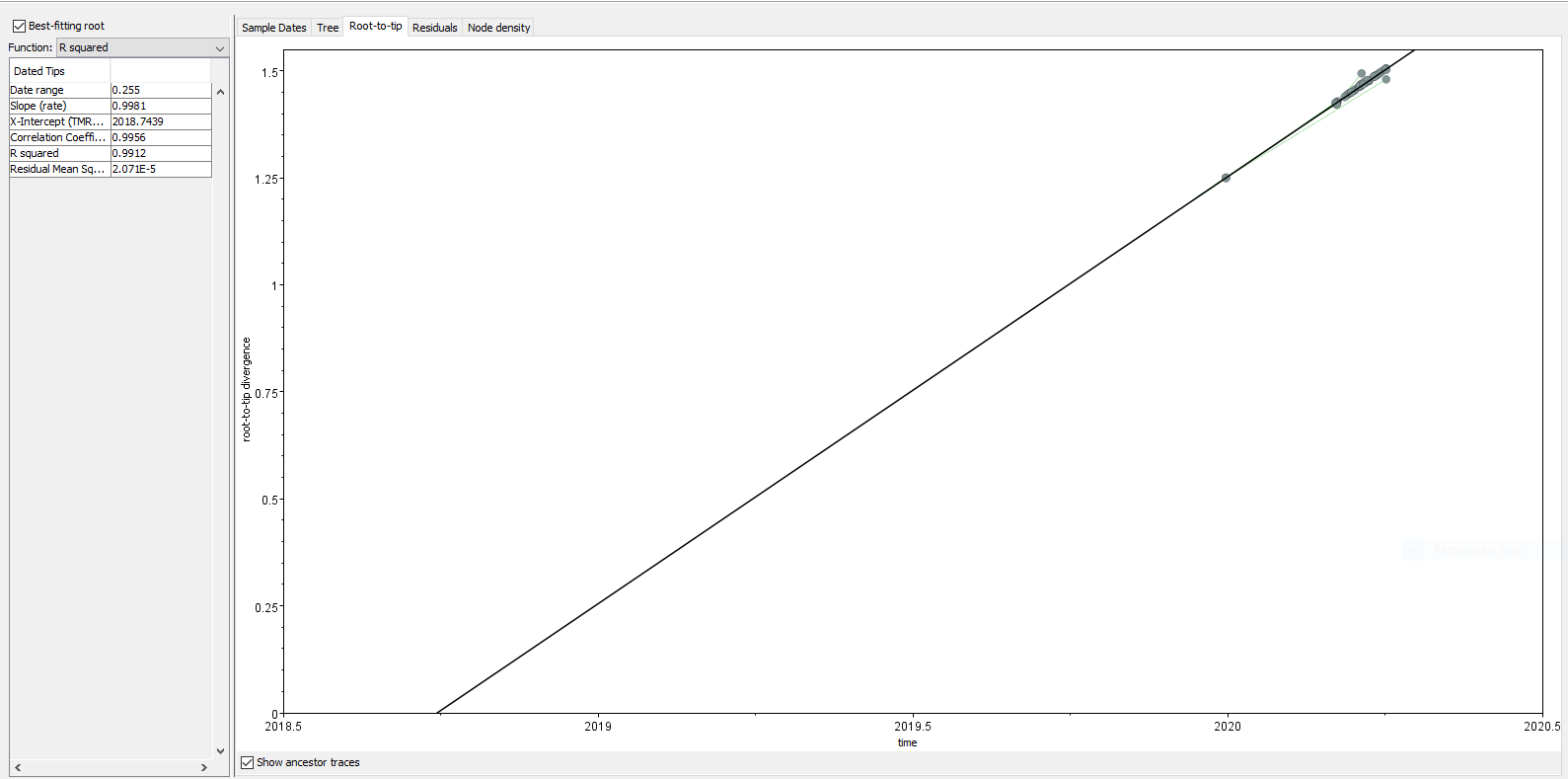
